# Integrin-identity shapes and mechanotemporally encodes the fibroblast transcriptome

**DOI:** 10.64898/2026.06.17.733008

**Authors:** Upnishad Sharma, Michele M. Nava, Luca F Witte, Nicholas Luginbühl, Huanding Ji, Michal Okoniewski, Rebecca Kottke, Barbara Treutlein, Reinhard Faessler, Daniel J Müller

## Abstract

Fibroblast transcriptomic states reflect physiological and pathological tissue contexts, yet the upstream determinants that stabilize these states remain poorly defined. Integrins mediate extracellular matrix (ECM) adhesion and biochemical signaling, but whether they encode mechanical constraints into stable transcriptomic programs is unclear. Using engineered mouse fibroblasts, bulk and single-cell transcriptomics, and controlled micromechanical confinement, we show that integrins can shape the transcriptomic landscape. The bulk transcriptome of fibroblasts expressing αV- and β1-class indicates a shared mechanosensitive baseline, except when these integrin classes are expressed individually. We also found integrin-specific gene clusters, including β1-class integrin-dependent enrichment of *Areg, Epha7, Lhpp* and *Igf2r*, which regulate development, regeneration and disease, and altered YAP1 targeted gene expression. At single-cell resolution under confinement, β1-class integrins sustain a progenitor-associated program, whereas their loss or αV-class enrichment promotes a constitutively activated state linked to injury repair and wound healing. Mechanical confinement and confinement duration further reshapes these states in an integrin-identity-dependent manner. Our findings establish integrin-identity as a determinant of how fibroblasts transduce mechanotemporal inputs from the cell surface to the nucleus.

## INTRODUCTION

Ubiquitously distributed across mammalian tissues, fibroblasts are principal architects of the extracellular matrix (ECM), coordinating its synthesis, assembly, crosslinking, and remodeling^1^. Through these activities, they define tissue architecture and mechanical integrity^2^, while regulating morphogenesis, stem cell niche maintenance^3^, and long-term organ homeostasis^1^. In addition to these functions, fibroblasts actively shape immune responses^4^, guide wound repair, and contribute to angiogenesis and epithelial–mesenchymal crosstalk^1,2,4,5,6^. Their functional breadth is further underscored in pathological settings, where dysregulated fibroblast activation drives fibrosis, tumor progression, cardiac remodeling, and chronic inflammatory diseases such as inflammatory bowel disease^1,2,4,5,6^. Collectively, these observations position fibroblasts not merely as structural support cells but as dynamic regulators of tissue state across development, homeostasis, and disease.

To execute these diverse roles, fibroblasts continuously integrate biochemical and biophysical information from their microenvironment^7^. Their immediate surroundings comprise a complex and evolving ECM, neighboring epithelial, endothelial, and immune cells, as well as gradients of soluble cytokines and growth factors^8^. Superimposed on this molecular landscape are mechanical cues, including matrix stiffness, viscoelasticity, tensile strain, shear stress, and spatial confinement^9,10,11,12,13^. Fibroblasts decode this multidimensional input through cell surface receptors, among which integrins constitute the principal molecular interface between ECM and cytoskeleton^14,15^. As obligate α/β-subunit heterodimers, integrins exhibit distinct ligand-binding specificities and cytoplasmic interaction profiles, enabling selective engagement with collagen, fibronectin, laminin, and other ECM components^16^. Through the assembly of focal adhesion complexes, integrins physically couple extracellular ligands to actin stress fibers, thereby transmitting mechanical forces across the plasma membrane and into the cytoskeletal network^9^.

Beyond facilitating adhesion, integrins function as bidirectional signaling hubs^16,17^. Outside-in signaling translates ECM composition and mechanical resistance into intracellular kinase cascades, Rho-family GTPase activation, and cytoskeletal reorganization, whereas inside-out signaling modulates integrin affinity and clustering in response to intracellular cues^18^. In fibroblasts, these processes regulate migration, contractility, matrix deposition, and myofibroblast differentiation^19^. Mechanical loading of integrin adhesions promotes force-dependent conformational changes in focal adhesion proteins, alters adhesion turnover, and modulates actomyosin tension^15^. Through these mechanisms, integrins actively shape tissue mechanics by adjusting ECM production and remodeling^14,20^. However, despite extensive characterization of integrin-mediated adhesion dynamics, their contribution to defining transcriptomic states remains insufficiently resolved.

Recent single-cell and spatial transcriptomic analyses have revealed substantial heterogeneity among fibroblasts across tissues and disease conditions, identifying distinct transcriptomic states associated with inflammatory^8^, myofibroblastic^2^, and tissue-supportive states^21^. These findings emphasize fibroblast plasticity and context dependence, yet the upstream determinants that specify fibroblast identity remain to be understood^6^. It is unclear whether the combinatorial expression of specific integrin heterodimers, which collectively constitute an “integrin-identity”, correlates with distinct transcriptomes. Because integrins differ in their ligand repertoire, force transmission, and recruitment of adaptor proteins such as talin, kindlin, and vinculin^22^, distinct integrin combinations could establish divergent mechanochemical signaling platforms^10,14,20,23,24,25,26^.

Emerging evidence positions mechanosensing and mechanotransduction as central regulators of fibroblast function in tissue-specific contexts^7,27,28,29,30^. Mechanical inputs, including substrate stiffness and geometric confinement, influence fibroblast proliferation, differentiation, and activation^9,12,13,31^. Force transmission from integrin-based adhesions to the nucleus occurs *via* the actin cytoskeleton and LINC (linker of nucleoskeleton and cytoskeleton) complex, enabling mechanical modulation of nuclear shape, chromatin accessibility, and transcription factor dynamics^32,33,34^. Mechanosensitive regulators such as YAP/TAZ, myocardin-related transcription factors (MRTFs), and other tension-responsive cofactors translate cytoskeletal tension into transcriptional outputs^35,36^. Yet, most mechanotransduction studies have treated focal adhesions as uniform signaling entities, without distinguishing the specific contribution of integrin heterodimers. Given that integrins exhibit distinct force-dependent behaviors and signaling outputs, it is plausible that integrin-identity selectively primes fibroblasts for particular mechanotranscriptomic responses.

This conceptual gap is significant. If the integrin composition determines how fibroblasts interpret and encode mechanical information at the level of chromatin and transcription, then the integrin-identity may represent a previously underappreciated axis of cell-state regulation. Such a mechanism could explain how fibroblasts acquire tissue-specific phenotypes in mechanically diverse environments, ranging from compliant lung parenchyma to rigid bone-associated connective tissues. It could also provide insight into how altered mechanics in fibrosis or tumor stroma reinforce pathological fibroblast states through integrin-dependent transcriptomic reprogramming. To test this, we set out to determine whether integrin-identity specifies the fibroblast transcriptome and shapes its response to mechanical input. Using a well-characterized mouse kidney-derived fibroblast line as a tractable and genetically manipulable model, we engineered cells with defined integrin-identities and combined bulk and single-cell transcriptomic profiling with controlled mechanical confinement. This design lets us first resolve that integrin composition sets the baseline transcriptome and then ask how it governs the transcriptomic response to mechanical constraint, linking integrin-identity, force transmission, and gene expression. Together, these experiments establish a conceptual framework in which integrin-identity modulates fibroblast transcriptomes in response to mechanical cues, providing a basis for understanding how integrin diversity can contribute to tissue specialization in homeostasis and to maladaptive remodeling in disease.

## RESULTS

### Cell surface integrin-identity determines steady-state transcriptome

To elucidate the extent to which the fibroblast state arises from the constitution of integrins and potentially transduces mechanical cues, we engineered isogenic mouse wild-type (WT) and pan-integrin-KO (pKO) fibroblasts. The latter were reconstituted with integrin cDNAs to derive fibroblasts that only differ in the expression of specific integrins on their surface, namely, pKO-αV/β1 fibroblasts, expressing heterodimers from β1- and αV-class integrins, pKO-β1 fibroblasts that express β1-class integrins only, and pKO-αV fibroblasts that express only αV-class integrins^26^. Furthermore, we derived from pKO-αV fibroblasts pKO-αVβ3 and pKO-αVβ5 fibroblasts that express only αVβ3 or αVβ5 integrins, respectively (Fig. 1a). The fibroblasts with different surface integrin-identities were cultured on fibronectin for 16 h and subjected to mRNA sequencing for bulk transcriptomic analysis. Principal component analysis showed that ∼60% of the transcriptomic variation across these fibroblasts was accounted for by the first three principal components, PC1, PC2, and PC3 (Fig. 1b,c; Supplementary Fig. 1). ∼26% of the transcriptome variation was attributed to PC1, causing WT and pKO-αV/β1 fibroblasts to cluster closer to the crossing of x- and y-axes, while pKO-β1 fibroblasts clustered towards the +x-axis, and pKO-αV, pKO-αVβ3, and pKO-αVβ5 fibroblasts, which exclusively express the αV-class integrins, clustered towards the -x-axis (Fig. 1b). This difference in clustering implied that PC1 corresponded to the axis that delineated the individual contributions of β1- and αV-class integrins to the varied gene expression. Moreover, ∼23% of the variation was attributed to PC2 (Fig. 1c), causing WT and pKO-αV/β1 fibroblasts to cluster together along the -x-axis, while pKO-β1 fibroblasts clustered towards the +x-axis, and pKO-αV, pKO-αVβ3, and pKO-αVβ5 fibroblasts, which express exclusively the αV-class integrins, spread along the y-axis (PC3). PC2 thus polarized the transcriptomic data of WT and pKO-αV/β1 fibroblasts that expressed the under-explored αVβ1 integrin heterodimer^14^, and of pKO-β1, pKO-αV, pKO-αVβ3, and pKO-αVβ5 fibroblasts that failed to express αVβ1 integrin. Furthermore, ∼11% of the transcriptome variation was attributed to PC3 (Fig. 1c), which primarily decoupled the overall gene expression variation within αV-class integrin expressing fibroblasts. The confirmation of the phenotypic classifications of PC1 and PC2 was supported by the list of top 30 most variable genes (PCA loadings) within the individual principal components PC1-PC4 (Supplementary Fig. 2). For instance, PC1 was dominated by the downregulation of *Itgav* and upregulation of *Itgb1*, PC2 was dominated by downregulation of *Itgb1*. In summary, integrin-identity, the combination of β1- and αV-classes of integrins and of αVβ3 and αVβ5 integrin heterodimers, shapes the steady-state transcriptome of fibroblasts. These findings highlight the effects that cellular integrin-identity takes in shaping the cellular transcriptome.

**Figure 1.**
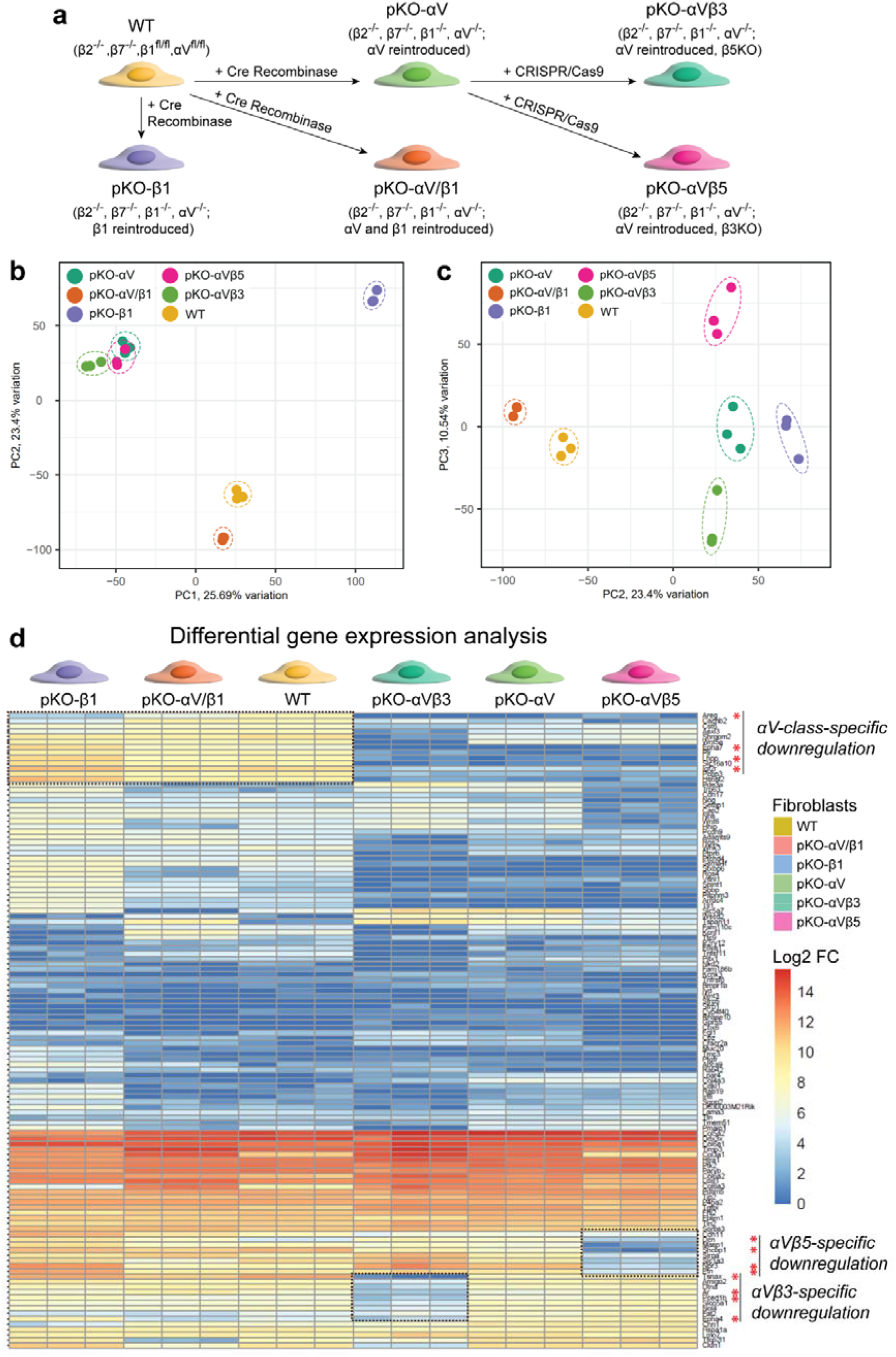
Integrin-identity determines the steady-state transcriptome of fibroblasts. **a**, Schematic of the genotype and engineering strategy of six different mouse fibroblasts showing different integrin-identities^26,24^. **b**, Principal component analysis (PCA) of the fibroblasts with principal components PC1 (x-axis) and PC2 (y-axis). **c**, PCA of the fibroblasts with PC2 (x-axis) and PC3 (y-axis). **d**, Differential gene expression analysis heatmap highlighting the upregulation or downregulation of particular sets of genes in an integrin class or integrin heterodimer specific manner. Red asterisks highlight genes that are upregulated in the β1-class or downregulated in an αV-class, αVβ3- or αVβ5-integrin specific manner. Color code of heatmap indicates log2 fold change (Log2 FC).

### Expression of specific sets of genes depends on integrin-identity

To further elucidate the effects of integrin-identity on the transcriptome, we performed differential gene expression analysis across the bulk transcriptome and observed entire clusters of genes to modulate their expression in an integrin-identity dependent manner (Fig. 1d). Genes like *Areg, Epha7, Lhpp*, and *Igf2r* exhibited upregulation in WT, pKO-αV/β1, and pKO-β1 fibroblasts that expressed β1-class integrins. This inference was corroborated by the downregulation of these genes upon loss of β1-class integrins in pKO-αV, pKO-αVβ3, and pKO-αVβ5 fibroblasts. The specific downregulation of genes like *Tsnax, Ar, Pced1b*, and *Epha4* was restricted to pKO-αVβ3 fibroblasts, which indicated the importance of both β1-class integrins and αVβ5 integrin for the baseline maintenance of these genes. Furthermore, downregulation of *Dcn, Npr3, Ptn*, and *Shcbp1* specifically in pKO-αVβ5 fibroblasts highlighted the dependence of these genes on the presence of both β1-class and αVβ3 integrins. Thus, the differential gene expression analysis delineates the dependence of several genes on either the integrin class or/and an integrin heterodimer.

### Integrin-identity-dependent transcription factors shape the transcriptome

The differential gene expression analysis identified the dependence of genes on specific integrin classes and integrin heterodimers. However, the mediator for these differential gene expressions remained unclear. To seek clarity on this issue, we explored the differentially expressed genes downstream of the various transcription factors (Fig. 2; Supplementary Fig. 3). Interestingly, most genes mentioned earlier, *Epha4, Epha7, Tsnax, Shcbp1, Ptn* and *Dcn*, showed a differential expression pattern downstream of YAP1. For example, *Epha7* was upregulated in pKO-β1 fibroblasts, while downregulated in all fibroblasts that lacked β1-class integrins. Additionally, we found that *Tsnax* and *Shcbp1* remained downregulated in an αVβ3 or αVβ5 integrin specific manner, respectively. Therefore, the YAP1-dependent regulation of several genes depended on the integrin-identity of the fibroblasts. This finding indicates the existence of a link between integrin-identity, integrin-mediated mechanotransduction, and integrin-dependent mechanoregulation of the cellular transcriptome.

**Figure 2.**
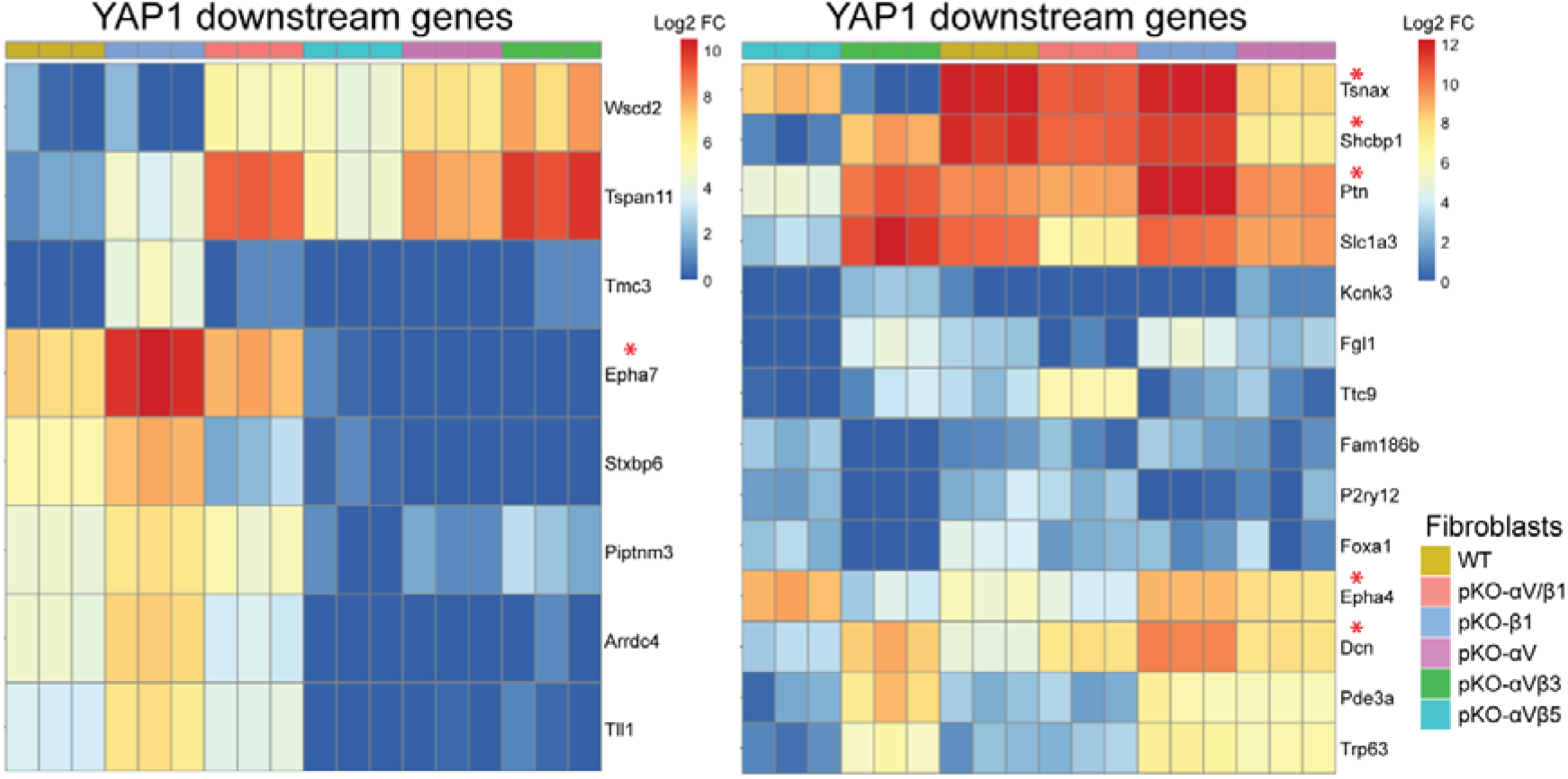
YAP1-regulated genes show integrin-identity dependent expression. The heatmap depicts upregulation or downregulation of genes downstream of the mechanosensitive transcription factor YAP1 in an integrin class or integrin heterodimer specific manner. Red asterisks, selected from Fig. 1d, highlight genes that are upregulated in an β1-class integrin dependent manner (*Epha7*) or downregulated in an integrin heterodimer-specific manner for αVβ3 integrin (*Tsnax*) or αVβ5 integrin (*Shcbp1*). Color code of heatmap indicates log2 fold change (Log2 FC).

### Mechanical confinement modulates transcriptomic output

To link integrin-identity with mechanotransduction, we examined how mechanical manipulation alters the single-cell transcriptome. Therefore, we combined static uniaxial confinement with live-cell imaging and sample harvesting for single-cell RNA sequencing (scRNA-seq) (Fig. 3a). Using this assay, we explored the effect of integrin-identity, confinement height and confinement duration on fibroblasts through the transcriptomic signatures obtained *via* scRNA-seq. We confirmed the effectiveness of the confinement assay on the fibroblasts by live-cell imaging where we used two confinement heights, 20 μm (control, unconfined) and 5 μm (confined). The 20 μm confinement height did not deform the fibroblast as indicated by the unchanged apical cellular and nuclear curvature in the x-z/y-z plane in the unconfined state (Fig. 3b). We confirmed this by atomic force microscopy (AFM)-based cell height measurements, which showed all fibroblast lines having comparable heights of ∼10 μm (Supplementary Fig. 4). However, upon confinement to 5 μm, the apical cellular and nuclear curvature of the fibroblasts flattened considerably in the x-z/y-z plane (Fig. 3b). To test whether the confinement itself and the duration of confinement influences the transcriptome, we confined (5 µm) the fibroblasts for 3 h and 16 h, before harvesting them for scRNA-seq (Fig. 3a), alongside with the unconfined (20 µm) control condition.

**Figure 3.**
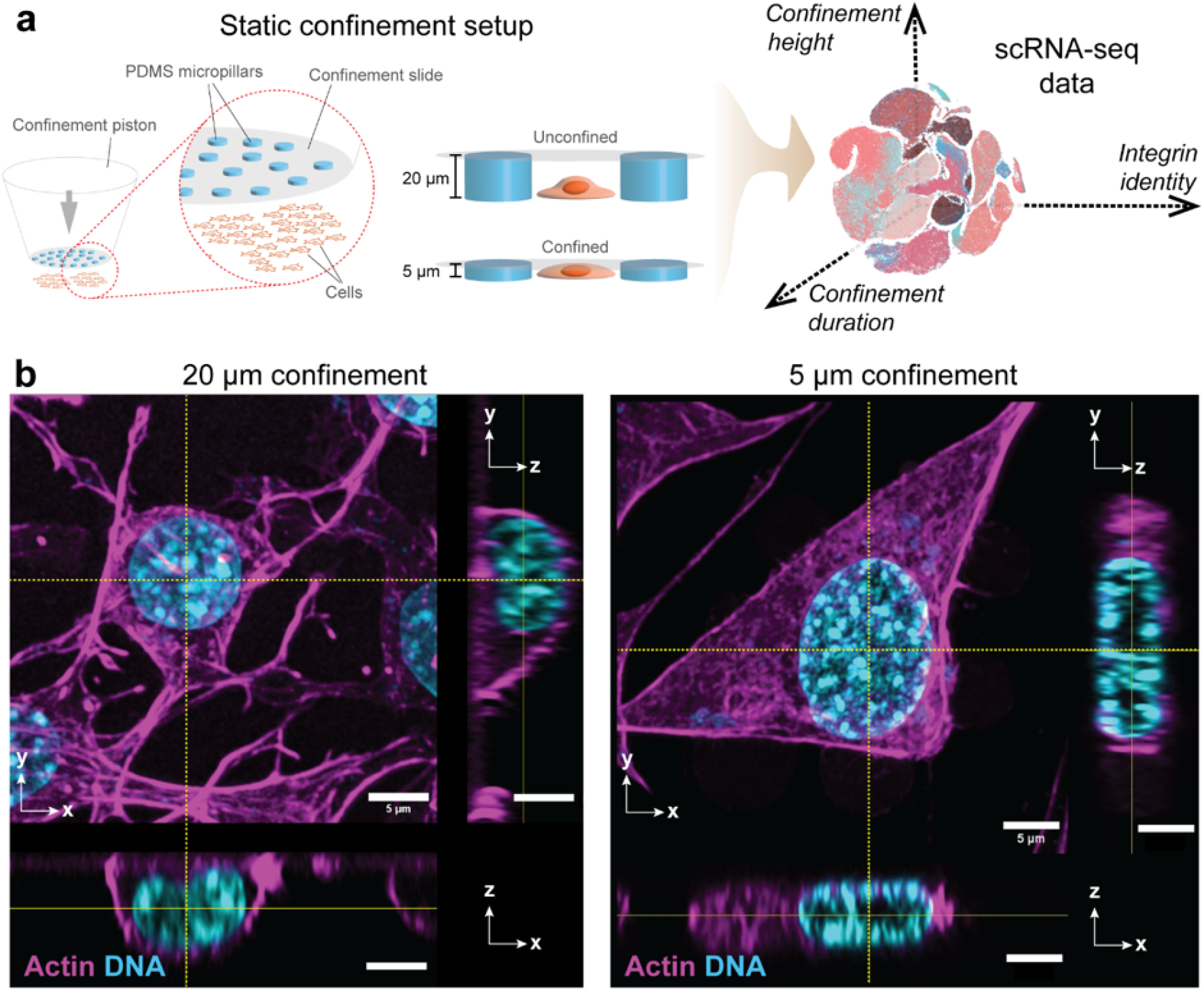
Basal transcriptomic output of fibroblasts is modulated both mechanically and temporally. **a**, Schematic of PDMS micropillars used to confine adherent fibroblasts to heights of 20 µm (control, unconfined) or 5 µm (confined) for 3 h or 16 h. After confinement, fibroblasts were subjected to scRNA-seq to characterize the transcriptome modulation along three parameters, integrin-identity, confinement height, and confinement time. **b**, Live-cell Airyscan confocal microscopy of fibroblasts during confinement. Actin (SiR-Actin-650™) and nuclear DNA (NucBlue™) are fluorescently labeled. Scale bar, 5 µm.

After 3 h of confinement, fibroblasts showed a distinctive clustering in the uniform manifold approximation and projection (UMAP) projections, which indicated that mechanical confinement influenced gene expression (Fig. 4a). This inference was further consolidated when fibroblasts confined for 16 h also exhibited different clustering in UMAP projections, though these clusters were less distinct than the ones obtained after 3 h (Fig. 4b). The weaker 16 h response may reflect adaptation, cytokine signaling, and migration during prolonged adhesion, which override the confinement signal. Thus, fibroblast gene expression at the single-cell level is modulated by confinement and temporally controlled.

### Integrin-identity influences the state and transcriptome of single fibroblasts

The confinement-dependent gene expression of fibroblasts became pronounced already when confined for 3 h. The gene expression across WT, pKO-αV/β1, and pKO-αV fibroblasts shows that integrin-identity alongside confinement modulate single-cell gene expression (Fig. 4a,b; Supplementary Fig. 5). Considerable changes in gene expression patterns over confinement time were observed across all three fibroblasts. Firstly, we observed that WT, pKO-αV/β1, and pKO-αV fibroblasts formed distinct clusters indicative of their integrin-identities. WT and pKO-αV/β1 fibroblasts clustered in an interlinked manner, whereas pKO-αV fibroblasts formed an separate cluster, which indicated the loss of β1-class integrins (Fig. 4b,c). Secondly, within the clusters marked for individual fibroblast types, we distinguished gene expression signatures specific to confinement times of 3 h and 16 h. These temporal changes in gene expression signatures markedly differed for WT, pKO-αV/β1, and pKO-αV fibroblasts. Lastly, the effect of confinement was most remarkable for pKO-αV/β1 fibroblasts for both confinement times and for WT fibroblasts at 16. On the contrary, pKO-αV fibroblasts minimally altered their gene expression signature in response to 3 h and 16 h long confinement. This muted response suggested that pKO-αV fibroblasts have a reduced capacity to diversify their state. To identify a specific mediator of this phenomenon, we analyzed the distribution of *Cd34* (Fig. 4c,d), a marker for inactivated fibroblast progenitors^6,21,37^, which enriched specifically in β1-class integrins in bulk mRNA analysis (Fig. 1d). pKO-αV fibroblasts completely lacked *Cd34* expression, compared to WT and pKO-αV/β1 fibroblasts, which implies that the presence of β1-class integrin primed fibroblasts to become activated, from an inactive precursor state (Fig. 4c,d). On the contrary, pKO-αV fibroblasts exhibited elevated levels of *Cxcl5*, which is an indicator of activated fibroblasts engaged in injury repair and wound healing^2^. These findings imply that the loss of β1-class integrins leads to a pre-activated state in fibroblasts and that the integrin-identity may indeed influence a role in triggering single-cell gene expression under mechanical confinement.

**Figure 4.**
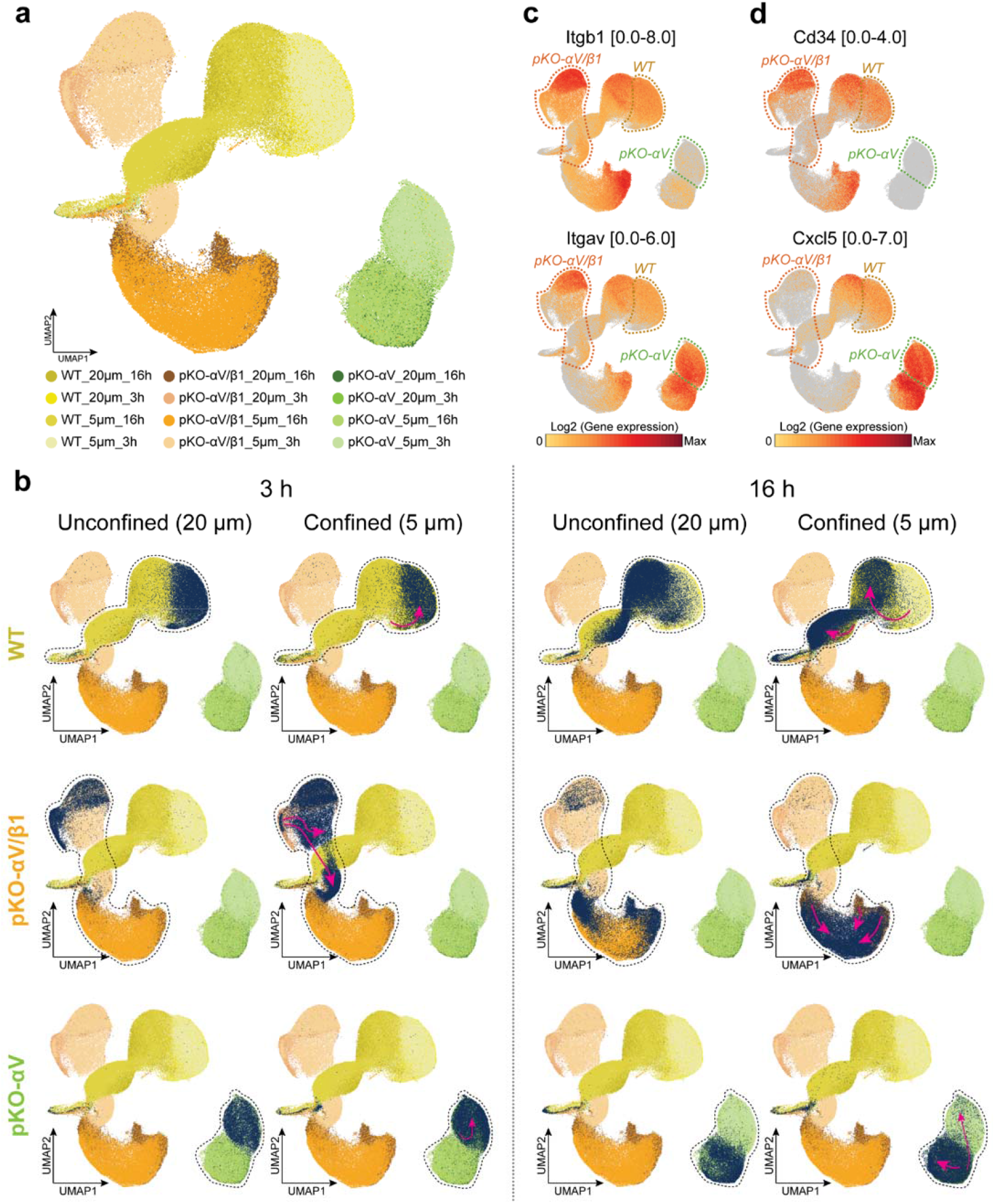
Integrin-identity defines the transcriptome response and state of confined fibroblasts. **a**, Uniform manifold approximation and projection (UMAP) of scRNA-seq from WT, pKO-αV/β1, and pKO-αV fibroblasts being unconfined (20 µm) and confined (5 µm) for 3 h or 16 h. Loss of β1-class integrins leads to an entirely separate clustering of fibroblasts. **b**, Comparative transcriptomic distribution (dark blue) of unconfined (20 µm) and confined (5 µm) fibroblasts for 3 h or 16 h. Magenta arrows indicate tentative trajectories of transcriptome modulation by confinement. Dotted outlines trace the transcriptomic expansion of WT, pKO-αV/β1, or pKO-αV fibroblasts. **c**, Expression profiles of *Itgb1* and *Itgav* across WT, pKO-αV/β1, or pKO-αV fibroblasts confined for 3 h. **d**, Expression profiles of *Cd34* and *Cxcl5* across WT, pKO-αV/β1, or pKO-αV fibroblasts confined for 3 h. Dotted lines in **c** and **d** mark regions specific to 3 h confinement. Color code of UMAPs in (**c**) and (**d**) indicates log2 of gene expression with quantitative range noted alongside the gene name.

### Integrin-identity differentially directs cellular response genes under confinement

Once confirmed that integrin-identity influences the gene expression alongside confinement height and time, we examined the genes differentially enriched in unconfined (20 μm) or confined (5 μm) WT, pKO-αV/β1, and pKO-αV fibroblasts after 3 h (Fig. 5). Unconfined WT fibroblasts enriched immune response-linked genes like *Cebpb* and *Ppp1r1b*, whereas confined WT fibroblasts enriched cell invasiveness and proliferation-linked genes like *Epha2, Klf9*, and *Gadd45a* (Fig. 5a). Unconfined pKO-αV/β1 fibroblasts enriched *Aqp5* and essential cytoskeletal component *Krt16*, while confined pKO-αV/β1 fibroblasts enriched *Pdgfb* and critical transcription factor genes like *Sox9, Cebpb, Nr4a1*, and *Nr4a2* (Fig. 5b). Unconfined pKO-αV fibroblasts enriched the cell-cell adhesion gene *Cdh6*, whereas confined pKO-αV fibroblasts enriched transcription factor genes like *Cebpb, Nr4a1*, and *Nr4a2*, similar to confined pKO-αV/β1 fibroblasts. Such enrichment indicates that the expression of these genes is shared among confined cells expressing αV-class integrins (Fig. 5c). We verified the distribution of these differentially expressed genes across WT, pKO-αV/β1, and pKO-αV fibroblasts using UMAPs (Fig. 5d). Thus, a considerable variation in the expression of genes related to cellular response, cell proliferation and transcription factors is induced by mechanical confinement, from which a subset of genes is expressed in an integrin-identity-dependent manner.

**Figure 5.**
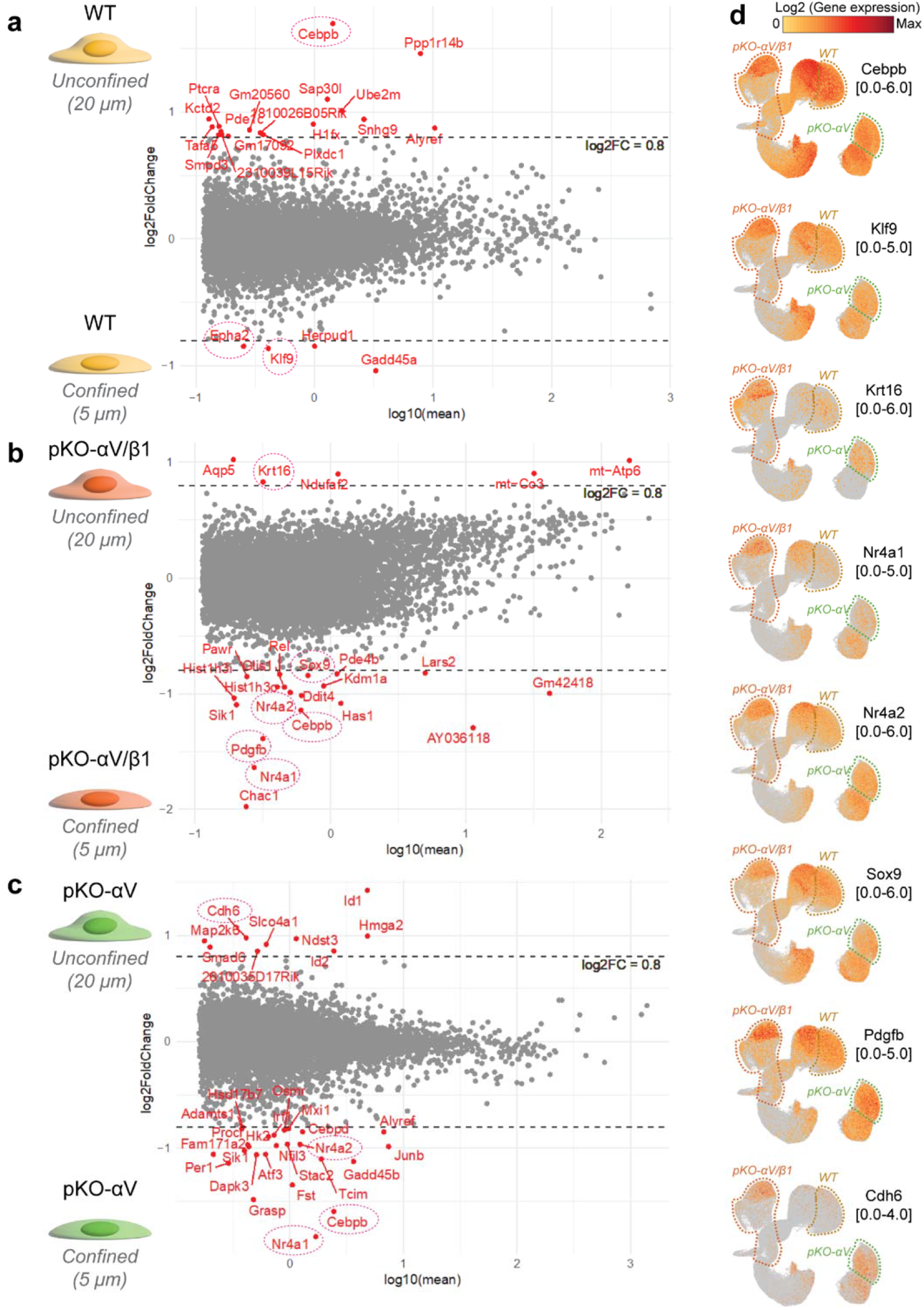
Integrin-identity differentially shapes confinement-induced gene expression. **a**, Differential gene expression analysis comparing gene enrichment in unconfined WT fibroblasts (+y-axis) relative to WT fibroblasts confined (-y-axis) for 3 h. **b**, Differential gene expression analysis comparing gene enrichment in unconfined pKO-αV/β1 fibroblasts (+y-axis) relative to pKO-αV/β1 fibroblasts confined (-y-axis) for 3 h. **c**, Differential gene expression analysis comparing relative gene enrichment in unconfined pKO-αV fibroblasts (+y-axis) to pKO-αV fibroblasts confined (-y-axis) for 3 h. Grey dashed lines in (**a**-**c**) mark the log2FC threshold chosen for relative gene enrichment and encircled (dotted red ellipses) mark key disease-associated genes. **d**, Confinement-induced expression variation of *Cebpb, Klf9, Krt16, Nr4a1, Nr4a2, Sox9, Pdgfb*, and *Cdh6* in UMAP projections of fibroblasts (dotted lines mark different fibroblasts lines confined for 3 h). Color code of UMAPs indicates log2 of gene expression with quantitative range noted alongside the gene name.

### Cytoplasm-nucleus mechanotransduction adopts distinct modalities

Given that the confinement induced gene expression depends on integrin-identity, we wondered what the mediators of this process could be. Earlier (Fig. 2), we observed several integrin-identity dependent genes downstream of the mechanosensitive transcription modulator YAP^35^. Therefore, we created YAP1-EGFP expressing WT, pKO-αV/β1, pKO-β1 and pKO-αV fibroblasts to morphologically identify the effects that confinement and integrin-identity could have on the relative cytoplasmic/nuclear localization of mechanosensitive YAP (Fig. 6a), independent of the morphological changes (area or aspect ratio) in the confined nucleus (Supplementary Fig. 6). For WT, pKO-αV/β1, pKO-β1 and pKO-αV fibroblasts, YAP localization to the nucleus increased upon confinement. In WT fibroblasts, confinement increased the nuclear recruitment of YAP by ∼22%, in pKO-αV/β1 fibroblasts by ∼67%, in pKO-β1 fibroblasts by ∼47% and in pKO-αV fibroblasts by ∼70% (Fig. 6b). While these results showed that confinement led to an increased relative nuclear recruitment of YAP^38^, they also showed that the degree of YAP recruitment was regulated by integrin-identity. To further explore the integrin-dependence of YAP nuclear localization, we immunostained WT, pKO-αV/β1, pKO-β1, and pKO-αV fibroblasts for the unphosphorylated active form of YAP (aYAP), which preferentially localized to the nucleus (Fig. 6c). All fibroblasts showed an aYAP nuclear/cytoplasmic ratio of >1.5, signifying the enhanced nuclear recruitment of aYAP, as expected (Fig. 6d), so that the effect of confinement on the aYAP amount was less pronounced than in YAP1-EGFP expressing fibroblasts. However, strikingly the aYAP nuclear/cytoplasmic ratio for pKO-β1 fibroblasts was ∼1.5 times higher than for WT, pKO-αV/β1, and pKO-αV fibroblasts (Fig. 6d). In summary, monitoring different modes of YAP signaling, either by total YAP or aYAP, highlights how mechanical confinement mechanotransduced to the nucleus differentially depends on the integrin-identity of fibroblasts.

**Figure 6.**
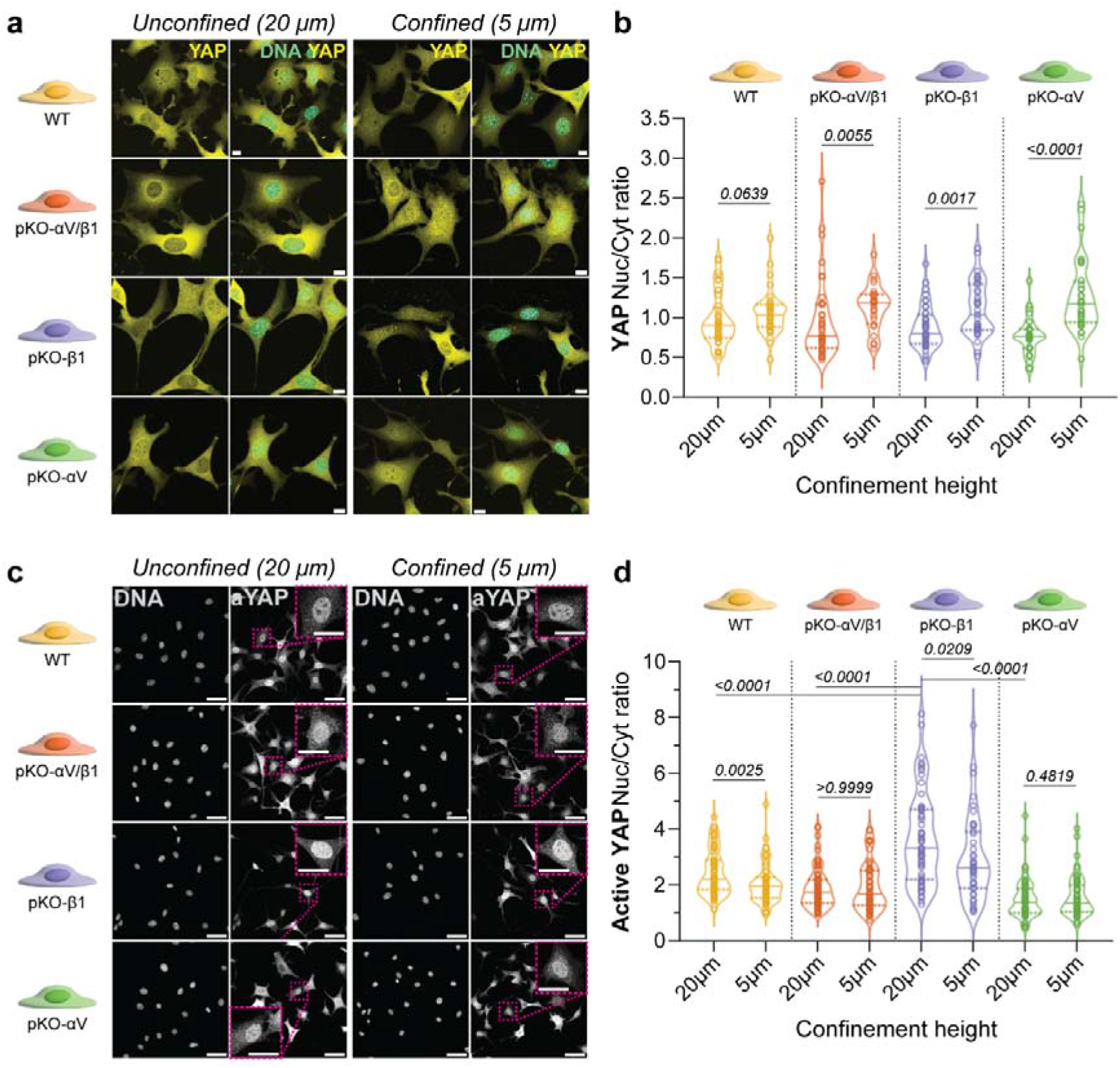
Integrin-identity sets distinct modes of confinement-induced YAP nuclear translocation. **a**, Confocal microscopy images of WT, pKO-αV/β1, pKO-β1, and pKO-αV fibroblasts expressing YAP1-EGFP (yellow) unconfined (20 µm) or confined (5 µm). Nuclear DNA has been labelled using NucBlue (cyan). Scale bars, 10 μm. **b**, Nuclear to cytoplasmic (Nuc/Cyt) ratio of YAP across WT [*n*(20 µm)= 41; *n*(5 µm)= 32], pKO-αV/β1 [*n*(20 µm)= 41; *n*(5 µm)= 24], pKO-β1 [*n*(20 µm)= 40; *n*(5 µm)= 35], and pKO-αV [*n*(20 µm)= 30; *n*(5 µm)= 32] fibroblasts. **c**, Active-YAP (aYAP) of immunolabelled WT, pKO-αV/β1, pKO-β1, and pKO-αV fibroblasts unconfined (20 µm) or confined (5 µm). Nuclear DNA has been labelled using NucBlue. Scale bars, 50 μm. (Inset) Zoom-in to better depict aYAP nuclear enrichment. Scale bars, 25 μm. **d**, Nuclear to cytoplasmic (Nuc/Cyt) ratio of aYAP across WT [*n*(20 µm)= 68; *n*(5 µm)= 61], pKO-αV/β1 [*n*(20 µm)= 79; *n*(5 µm)= 57], pKO-β1 [*n*(20 µm)= 56; *n*(5 µm)= 40], and pKO-αV [*n*(20 µm)= 74; *n*(5 µm)= 61] fibroblasts. In violin plots (**b** and **d**), each data point corresponds to a single cell. Solid line represents the median, the dotted line indicates the interquartile range. The width of the violin plot represents the density of data points at that value with wider sections indicating higher density. *p*-values obtained from unpaired, two-sided, Mann-Whitney U-test. *p*-values that compare unconfined (20 µm) pKO-β1 fibroblasts to unconfined (20 µm) WT, pKO-αV/β1 and pKO-αV fibroblasts were obtained using a two-way ANOVA with multi-comparison Tukey’s test.

## DISCUSSION

The transcriptomic states of fibroblasts closely mirror physiological and pathological conditions^1,2,4,5,6^. Yet the upstream determinants that evoke and stabilize these states remain poorly defined. Biochemical features of the ECM, sensed primarily through integrins, have been established as regulators of cell-fate decisions during development and as drivers of maladaptive fibroblast activation in fibrosis, tumor progression, and organ failure^39,40^. However, it remains unclear whether integrins also function as central mediators that translate mechanical constraints into context-specific transcriptomic programs^21,41,42^. Here, we show that fibroblasts, depending on their integrin-identity, mount distinct transcriptomic responses to mechanical inputs. By establishing a relationship between integrin-identity and mechanosensitive gene regulation, our work advances the understanding of the cell surface– to–nucleus axis of mechanotransduction and its contribution to fibroblast state specification.

Our results demonstrate that defined integrin-classes modulate the global transcriptomic landscape of fibroblasts. Principal component analysis reveals a highly similar bulk transcriptome in WT and pKO-αV/β1 fibroblasts as they cluster closely across multiple components (Fig. 1b). This convergence suggests that preserved expression of αV- and β1-class integrins together contribute substantially to maintaining a shared transcriptomic baseline. Differential expression analysis identifies discrete gene clusters that are selectively regulated by specific integrin classes or individual heterodimers (Fig. 1d). Notably, we detect altered expression of genes downstream of YAP1, a central mechanosensitive transcription regulator, in an integrin-dependent manner (Fig. 2). This observation implicates integrin composition in shaping mechanotransductive signaling outputs at the level of transcription. Because these YAP1 target genes were assigned from curated gene sets and YAP activity itself was not manipulated, YAP1 is best regarded here as a candidate mediator; establishing its causal contribution will require integrin-specific loss- and gain-of-function perturbation of YAP signaling. However, delineation of the complete gene network linking distinct integrin heterodimers to YAP1 activity and broader transcriptomic networks will require further mechanistic dissection. Collectively, the observed correspondence between integrin class composition, heterodimer expression, and transcriptomic similarity supports our central hypothesis that cellular integrin-identity is tightly coupled to fibroblast transcriptomic state and thereby contributes to defining the cellular identity.

Furthermore, we identify discrete gene clusters that are selectively modulated according to integrin class and heterodimer composition. Differential expression analysis reveals that a subset of genes is specifically dependent on β1-class integrins for sustained expression. These β1⍰dependent effects could alternatively reflect an indirect role of the absence of an αV⍰integrin–mediated inhibitory signal, rather than a strictly direct activating role for β1⍰class integrins. Among these, *Areg, Epha7, Lhpp*, and *Igf2r* are consistently reduced in fibroblasts lacking β1-class integrins, indicating a non-redundant regulatory contribution of this integrin class (Fig. 1d). *Areg* (amphiregulin) encodes an autocrine growth factor implicated in tissue repair and fibroblast activation. *Epha7*, a member of the ephrin receptor family, functions as a key regulator of morphogenetic processes and tissue patterning^43^. *Lhpp* encodes a histidine phosphatase that has recently been characterized as a tumor suppressor with roles in cellular growth control and metabolic regulation^44^. *Igf2r*, the insulin-like growth factor 2 receptor, is an imprinted gene with established functions in growth regulation and developmental signaling^45^. The coordinated dependence of these functionally diverse genes on β1-class integrins underscores the breadth of transcripts influenced by integrin composition. Taken together, these findings extend the established role of integrins beyond adhesion and cytoskeletal organization, positioning β1-class integrins as upstream regulators of gene networks central to development, growth control, and disease-associated processes. This integrin-dependent transcriptomic specificity reinforces the concept that integrin-identity contributes directly to shaping fibroblast functional state.

We also demonstrate that both the mechanical confinement and its duration modulate gene expression at single-cell resolution in an integrin-dependent manner. These findings extend the integrin repertoire beyond adhesion, implicating integrin-identity in how fibroblasts transcriptomically encode mechanical constraints. Single-cell transcriptomic profiling of the distinct fibroblast lines reveals that β1-class integrins are required to maintain a fibroblast progenitor transcriptomic program. This state appears to preserve cellular plasticity, enabling subsequent activation into myofibroblasts or differentiation into specialized fibroblast subtypes in response to contextual cues^46^. In contrast, fibroblasts lacking β1-class integrins, or those relatively enriched in αV-class integrins, display elevated expression of *Cxcl5*, indicative of a constitutively activated fibroblast transcriptomic state (Fig. 4d)^46^. Thus, integrin composition not only influences baseline gene expression profiles, but also predisposes fibroblasts toward distinct functional phenotypes. Mechanical confinement further modifies these transcriptomic states. Both short-term and prolonged confinement induce measurable transcriptomic changes. This observation indicates that fibroblasts integrate mechanical signals cumulatively, with sustained confinement reinforcing or reshaping transcriptomic programs over time^29,47^. Moreover, pathway-level analyses reveal that nuclear response programs and inflammatory signaling pathways are selectively inducible by mechanical constraint in an integrin class–specific manner (Fig. 5d). For example, mechanical confinement triggers the expression of genes related to cellular responses, such as proliferation, invasiveness, immune response, wound healing, cytoskeleton modification, and transcription factors. Thus integrin-identity governs the sensitivity and quality of mechanotranscriptomic responses that are important for physiological and pathophysiological behavior. We note that the present study is conceptual in scope: it establishes that integrin-identity and mechanical confinement jointly shape the fibroblast transcriptome, rather than providing a systematic characterization of how graded confinement heights, durations, and additional mechanical parameters quantitatively tune this response. Such parametric dissection, enabled by the framework introduced here, is an important direction for future work. In summary, our findings establish integrin composition as a key determinant of how fibroblasts interpret mechanical confinement to define and stabilize transcriptomic cell states (Fig. 7).

**Figure 7.**
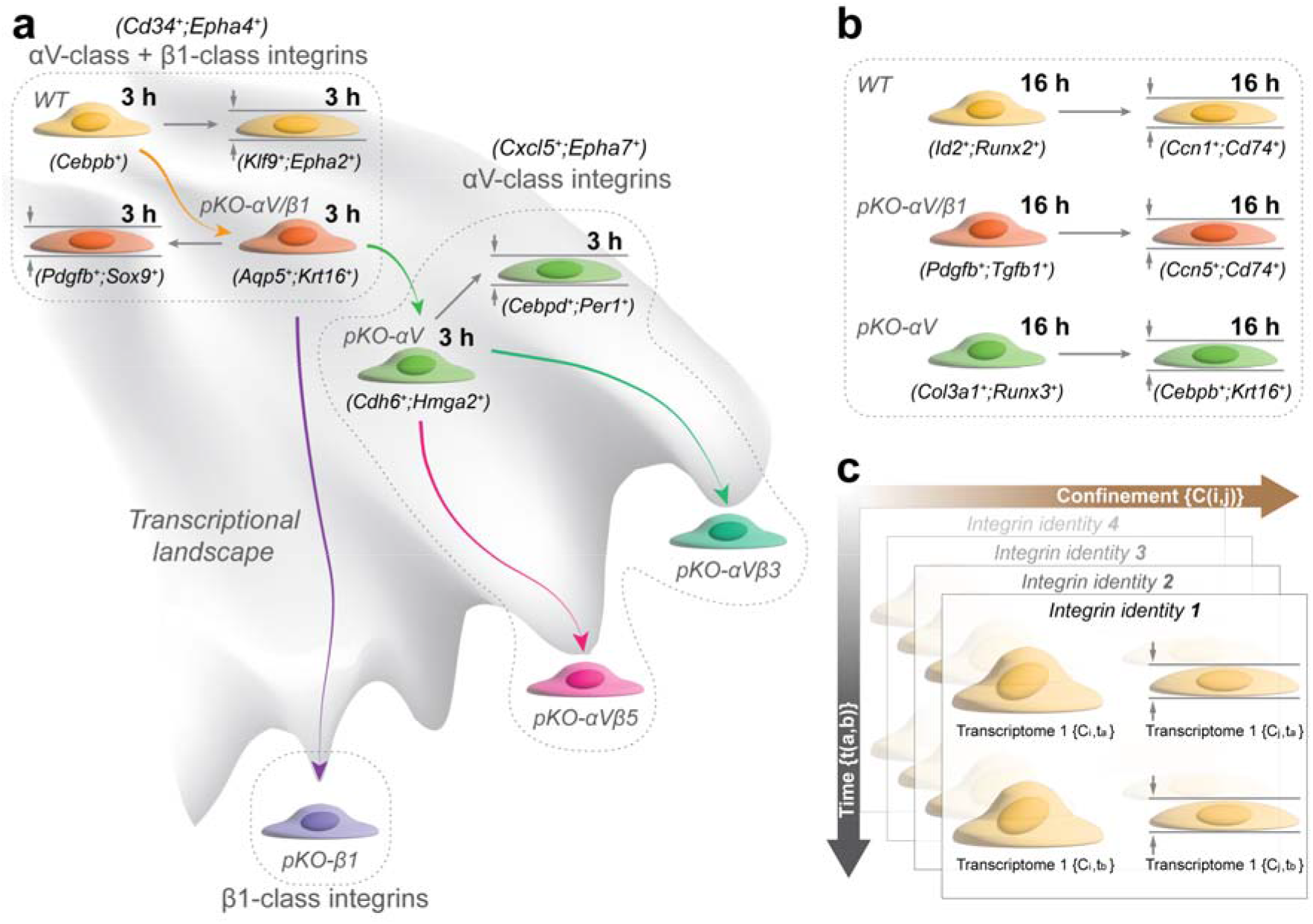
Integrin-identity shapes the fibroblast transcriptomic landscape depending on mechanical confinement and confinement time. **a**, Schematic summarizing how integrin-identity and mechanical confinement shape the transcriptome of fibroblasts adhering to fibronectin in the unconfined (20 µm) and confined (5 µm) states for 3 h. Expression of divergent integrin classes and/or heterodimers generates distinct transcriptomic identities, with upregulation of the indicated genes specific to particular integrin classes; mechanical confinement further alters the transcriptome, as resolved by single-cell RNA sequencing. Main identifying genes for the unconfined and confined states are shown in brackets. **b**, As in (**a**), for fibroblasts adhering to fibronectin in the unconfined (20 µm) and confined (5 µm) states for the extended duration of 16 h, showing that prolonged confinement further reshapes the transcriptome in an integrin-class–specific manner. **c**, Summary schematic integrating (**a**) and (**b**): integrin-identity sets the baseline transcriptomic state of the fibroblasts, while mechanical confinement and time, drive integrin-specific trajectories across the transcriptomic landscape. Indicated are different integrin identities shifting the transcriptome along the confinement and confinement time axis.

Finally, our data provide evidence that distinct integrin classes are required to establish and maintain discrete functional states in fibroblasts exposed to mechanical confinement. This requirement offers a mechanistic explanation for the diversity of integrin expression observed across tissues and disease contexts, linking integrin heterodimer composition to the functional versatility of fibroblasts during homeostasis and pathological remodeling. Perturbations in integrin balance, whether through loss of specific integrins or enrichment of others, may therefore shift fibroblasts into alternative transcriptomic states. Such shifts are likely to arise from altered cell surface–to–nucleus mechanotransduction^35,36^, resulting in constrained or maladaptive gene expression programs. Consequently, dysregulated integrin-identity may represent a critical upstream determinant of aberrant fibroblast activation in disease.

## METHODS

### Fibroblasts culture and maintenance

Mouse pKO-αV, pKO-β1 and pKO-αV/β1 fibroblast lines were generated from fibroblasts (floxed parental) derived from the kidney of 21-day-old male mice carrying floxed αV and β1 alleles (αV^flox/flox^, β1^flox/flox^), and constitutive β2 and β7 null alleles (β2^−/−^, β7^−/−48^). Individual kidney fibroblast clones were immortalized by retroviral delivery of the SV40 large T. Then, the floxed and immortalized fibroblasts were retrovirally transduced with mouse αV and/or β1 integrin cDNAs while the endogenous floxed β1 and αV integrin loci were deleted by adenoviral transduction of *Cre* recombinase. Fibroblasts were cultured in high⍰glucose DMEM (Gibco, #10566016) supplemented with 10% v/v heat⍰inactivated fetal bovine serum (FBS; Invitrogen, #10082147), and 1% v/v penicillin–streptomycin (Sigma, #P4333). Cultures were maintained at 37□°C in a humidified 5% CO_2_ incubator and passaged at 70-80% confluence using 0.25% v/v trypsin–EDTA (Gibco, #25200072). Fibroblasts were used between passages P10–P20 to minimize senescence⍰associated changes. Mycoplasma testing was performed regularly using a PCR⍰based kit (Merck, #MP0035)

### Bulk RNA sequencing of fibroblasts

#### RNA isolation for bulk RNA sequencing

For bulk transcriptomics experiments, cells were cultured for 16 h in 6-well cell culture plates (Costar) prior to harvesting. 10^6^ cells were detached using TrypLE Express (Gibco, #12604013) at 37°C for two minutes. Cell disruption was performed using QIAshredder homogenizers (Qiagen, #79654) according to the manufacturer’s protocols. The RNeasy Mini Kit (Qiagen, #74104) was used to extract RNA, following the manufacturer’s instructions and including the following optional steps from the protocol: - addition of β-mercaptoethanol (Sigma-Aldrich, #M6250) to buffer RLT to denature endogenous RNAses - additional centrifugation for elimination of Buffer RPE carryover – addition of 30 units RNase-Free DNase (Qiagen, #79254) to isolated RNA samples (instead of performing on-column digestion)

#### -mRNA enrichment and next generation sequencing

All steps after RNA isolation were performed by the Genomics Facility Basel. Sample integrity was confirmed using RNA ScreenTape on a TapeStation (Agilent, #5067⍰5576). Libraries were prepared from 200 ng of total RNA, using TruSeq stranded mRNA reagents (Illumina, #20020594) according to the vendor’s guideline. SPRI beads (Beckman Coulter, #B23318) were used after library pooling to deplete adapter dimers. Before and after SPRI treatment, library quality was confirmed using a Fragment analyzer (Agilent, #DNF⍰473⍰0500). Obtained samples were sequenced on NovaSeq SP flow cell (Illumina, #20027464) in single-end mode and a read length of 101 bases.

### Confinement setup

Mechanical confinement was achieved using PDMS⍰based microfabricated static confiner devices (4D Cell, #CSOW 620). 6-well cell culture dishes (Mattek, #P06G-1.5-20-F) were coated with 50 µg ml^−1^ bovine plasma fibronectin (341631, Calbiochem, Merck Millipore) for 1 h at 37°C. The coverslips with 20 and 5 µm PDMS micropillars were coated with 2% w/v bovine serum albumin in 1X PBS and incubated for 1 h at 37°C. 7x10^4^ fibroblasts were seeded and allowed to adhere for 1 h at 37°C before confinement was applied by placing the PDMS lid onto the fibroblasts.

### Single-cell RNA sequencing

Single⍰cell suspensions were generated by enzymatic dissociation using Accutase (Sigma-Aldrich, #A6964) for 10 min at 37 °C, followed by filtration through a 40 µm strainer. Viability (>80%) was confirmed using trypan blue. Treated samples were processed by the Genomics Facility Basel using the Chromium Next GEM Single Cell 3’ HT Reagent Kits v3.1 (Dual Index, #1000492) and ∼2x10^4^ cells were loaded per channel of a droplet⍰based microfluidic system of a Chromium X device (10x Genomics, #1000464). Samples were sequenced to a depth of 2x10^4^ reads per cell on 2 NovaSeq S4 lanes (Illumina, # 20027466) after confirming library quality via a Fragment Analyzer (Agilent, #DNF⍰473⍰0500)

### Imaging of fibroblasts under confinement

Live imaging was performed using an inverted Airyscan confocal microscope (LSM-980, Zeiss) equipped with an environmental chamber (37□°C, 5% CO_2_). Fibroblasts were imaged in DMEM supplemented with 10% v/v FBS and 1% v/v penicillin–streptomycin (Sigma, Cat. P4333). Time⍰lapse imaging was conducted at 1 min intervals for up to 10 min using a 40× water⍰immersion objective (Zeiss) using the SR-4Y multiplex mode. Laser power and exposure times were optimized to minimize phototoxicity. Z⍰stacks were acquired when required and processed into maximum⍰intensity projections.

### Generation of stable GFP-labelled mechanosensor expressing cell lines

Mechanosensor constructs (GFP⍰tagged human YAP1) were cloned into a Sleeping Beauty (Addgene, #60523) backbone under a constitutive bidirectional promoter. Following antibiotic selection using 2 µg ml^−1^ puromycin, GFP positive cells were enriched by FACS sorting based on GFP intensity (Sony MA900). Enriched cell lines were validated by fluorescence microscopy and functional assays assessing mechanosensor responsiveness.

### Immunostaining for active YAP

After the confinement and live-imaging, as described above, confinement was released and the cells were fixed in 4□% v/v paraformaldehyde (Electron Microscopy Sciences, #15710) for 15□min at room temperature, followed by three washes in phosphate-buffered saline (PBS; Gibco, #10010-023). Permeabilization was performed with 0.1□% v/v Triton□X-100 (Sigma-Aldrich, #T8787) in PBS for 10□min, and non-specific binding was blocked with 5% w/v bovine serum albumin (BSA; Sigma-Aldrich, #A9647) in PBS for 1□h at room temperature.

Cells were incubated overnight at 4°C with rabbit anti-YAP (non-phospho-Ser127) antibody (Cell Signaling Technology, #4491; 1:200 dilution in 1% w/v BSA in PBS). After washing three times in PBS, samples were incubated for 1□h at room temperature with Alexa□Fluor□568-conjugated goat anti-rabbit IgG secondary antibody (Invitrogen, #A-11011; 1:500 dilution in 1□% w/v BSA in PBS). Nuclei were counterstained with DAPI (ThermoLFisher Scientific, #D1306; 300□nM in PBS for 5□min). Immunostained cells were imaged using a Leica Falcon SP8 confocal microscope equipped with a 40× oil-immersion objective.

Fluorescence intensity and subcellular localization of active YAP were quantified using FIJI with identical acquisition and analysis settings across all samples.

### Data analysis of sequencing data

*Bulk RNA sequencing data*. For the transcriptomic sequencing data, adapter sequences were trimmed off using trimmomatic^49^. Leading and trailing bases were cut off with an average quality score below 3, inner bases were cut off at an average quality below 15 in a 4 bp sliding window. Trimmed reads were aligned to the mouse reference GRCm39^50^ using HISAT2^51^. Samtools^52^ was used for sorting and indexing the obtained reads. For alignment of RNA sequences, StringTie^53^ was used. Subreads featureCounts^54^ was used to obtain feature counts of the identified transcripts. Count tables were exported in csv format for further analysis. The integrative genomics viewer was used for visualization of transcriptomic data. Analysis of the transcriptomic data was conducted in R unless mentioned otherwise. Differential gene expression analysis was conducted on the mean per-gene read counts of three replicates for each cell line. The edgeR package was used to fit a negative binomial generalized log-linear model and conduct likelihood ratio tests on the model coefficients. For all further analyses, only genes with a false discovery rate <0.01 were used. Data exploration and visualization includes dimensionality reduction via principal component analysis (PCA) and plotting differential gene expression for relevant sets of genes. A primary investigation of genes of interest was conducted via the GeneCards Suite website^55^, for relevant hits, primary sources were consulted and are cited in the text.

*Single*⍰*cell RNA*⍰*sequencing data*. Raw data were processed using Cell Ranger v7.0.1. Downstream analysis was performed in Seurat (version 5.1). Cells with >15% mitochondrial reads, <200 detected genes, or extreme UMI counts were excluded. Data were normalized using SCTransform, and dimensionality reduction was performed using PCA followed by tSNE and UMAP. Clusters were identified using a shared nearest⍰neighbor graph and annotated using canonical mouse fibroblast markers (*e*.*g., Vim, Col1a1, Fn1, Acta2*). Differential expression was assessed using Wilcoxon rank⍰sum tests. A primary investigation was conducted in the 10x Genomics Loupe Browser v6.5.0. Dimensionality reductions via UMAP and t-SNE were investigated for different combinations of datasets to identify potential clusters manually and via graph-based clustering. Enriched cluster motifs were investigated to identify transcriptomic signatures of identified clusters.

### Image analysis

Image processing was performed using FIJI (ImageJ). Background subtraction, drift correction, and segmentation were applied as appropriate. Nuclear and cytoplasmic regions were segmented using thresholding. Mechanosensor activity (GFP intensity ratios) was quantified along defined regions of interest inside and outside of the nucleus as previously described^56^.

### Statistical analysis

Statistical analyses were performed using GraphPad Prism (version 10.1). Comparisons between two groups were assessed using a two⍰tailed unpaired Mann-Whitney U-test. Multi-group comparisons of nuclear YAP localization between unconfined (20 µm) pKO-β1 fibroblasts and unconfined WT, pKO-αV/β1, and pKO-αV fibroblasts (Fig. 6d) were performed using a two-way ANOVA followed by Tukey’s multiple-comparison test. Unless otherwise stated, data are presented as median ± 95% confidence interval. Statistical significance was defined as *p* < 0.05.

## ACKNOWLEDGEMENTS

We thank Single Cell Facility at DBSSE, ETH Zurich, for support with confocal microscopy. We thank M. Feldkamp and C. Beisel of the Genomics Facility Basel for the single and bulk RNA sequencing experiments. We thank all the members of the biophysics laboratory for critical and constructive discussions during the initial phase of the project. We further thank H.J. Kaltenbach and J. Stelling for helpful statistical and conceptual discussions. This work was co-financed by the Swiss National Science Foundation (SNSF Grant No. 31003A-182587) and the NCCR Molecular Systems Engineering (SNSF Grant No. 51NF40-205608).

## AUTHOR CONTRIBUTIONS

U.S. and D.J.M. designed the experiments and discussed the project with R.F. and M.M.N.. U.S. performed cell confinement experiments with L.F.W. and M.M.N.. L.F.W. and M.O. analyzed the bulk and single-cell RNA sequencing. M.O., H.J. and N.L. further performed quality check and analysis on the single-cell RNA seq data. U.S. and M.M.N. performed confocal imaging, and analyzed the experimental data. R.K. and U.S. together generated the YAP-GFP expressing stable fibroblast lines. U.S. and D.J.M. wrote the first draft of the manuscript. All authors discussed the experiments and analysis and reviewed the manuscript.

