## Supplementary figures for "Integrin-identity shapes and mechanotemporally encodes the fibroblast transcriptome"

**Supplementary Information**

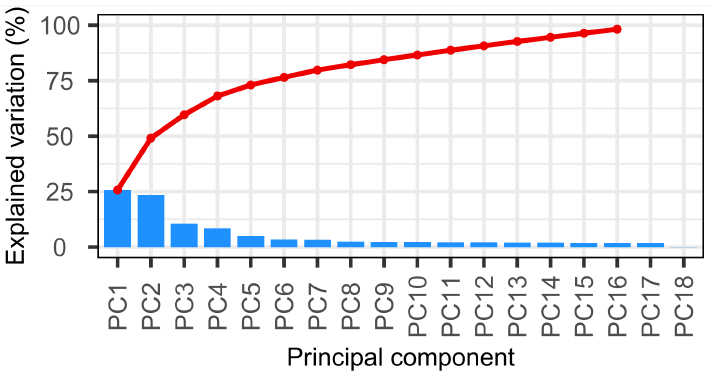

**Supplementary Figure 1. The first 18 principal components (PCs) depict the number of PCs required to explain percentage variability of the bulk transcriptomic data.** Elbow and bar plot depicts the contributions from individual PCs (blue bars) alongside the cumulative variation (red) across the PCs. The figure relates to Fig. 1b, which shows plots for the first three PCs, PC1-PC3.

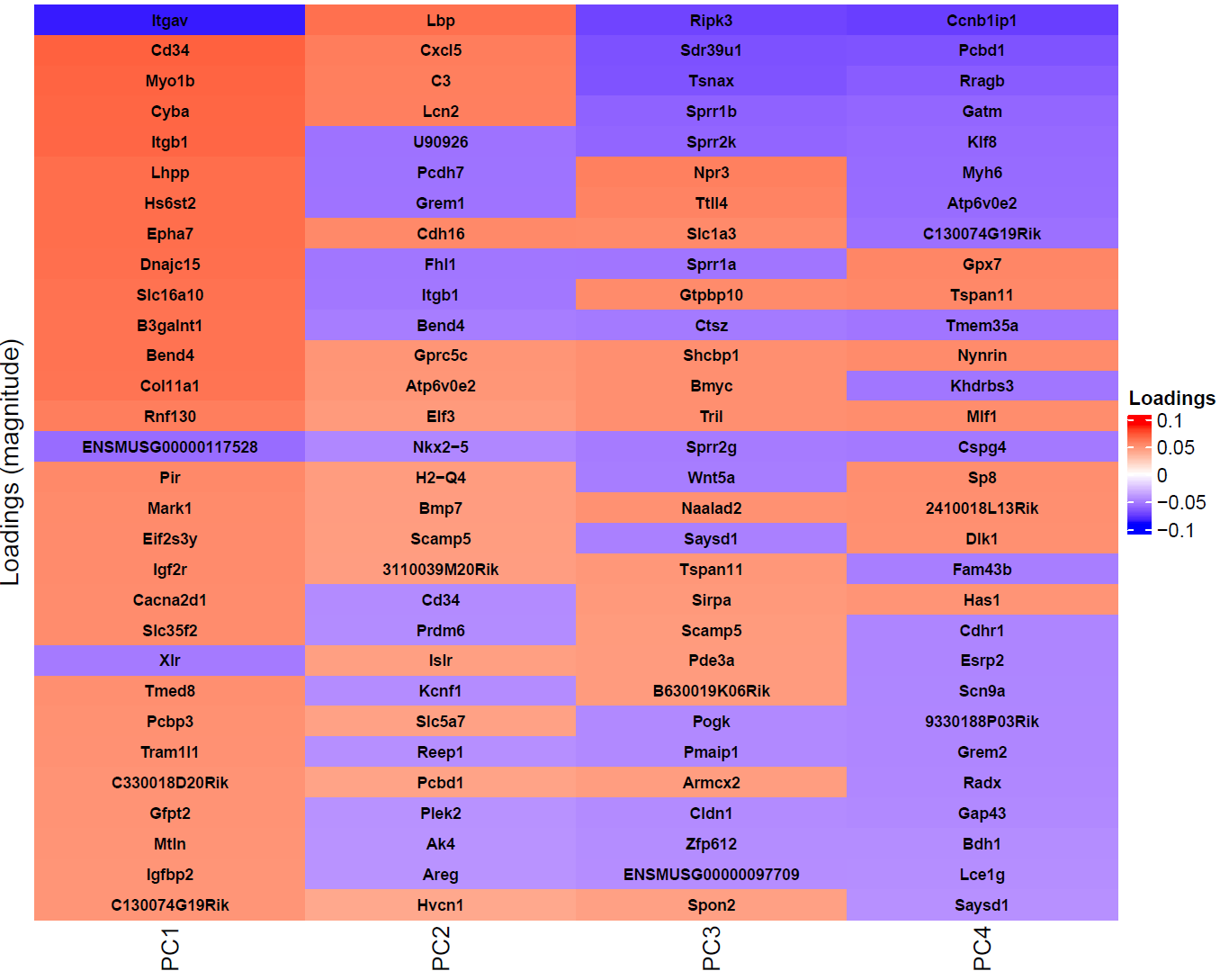

**Supplementary Figure 2. Top 30 genes that dominate the variability of the principal components PC1–PC4 in the bulk transcriptomic data.** The values are color-coded, with positive values depicted in red and negative values in blue. These loadings identify which variables have the strongest influence on each principal component. For targets which could not be successfully mapped to genes, only Ensembl IDs are provided. Color bar quantifies the loadings as indicated. The figure relates to Fig. 1.

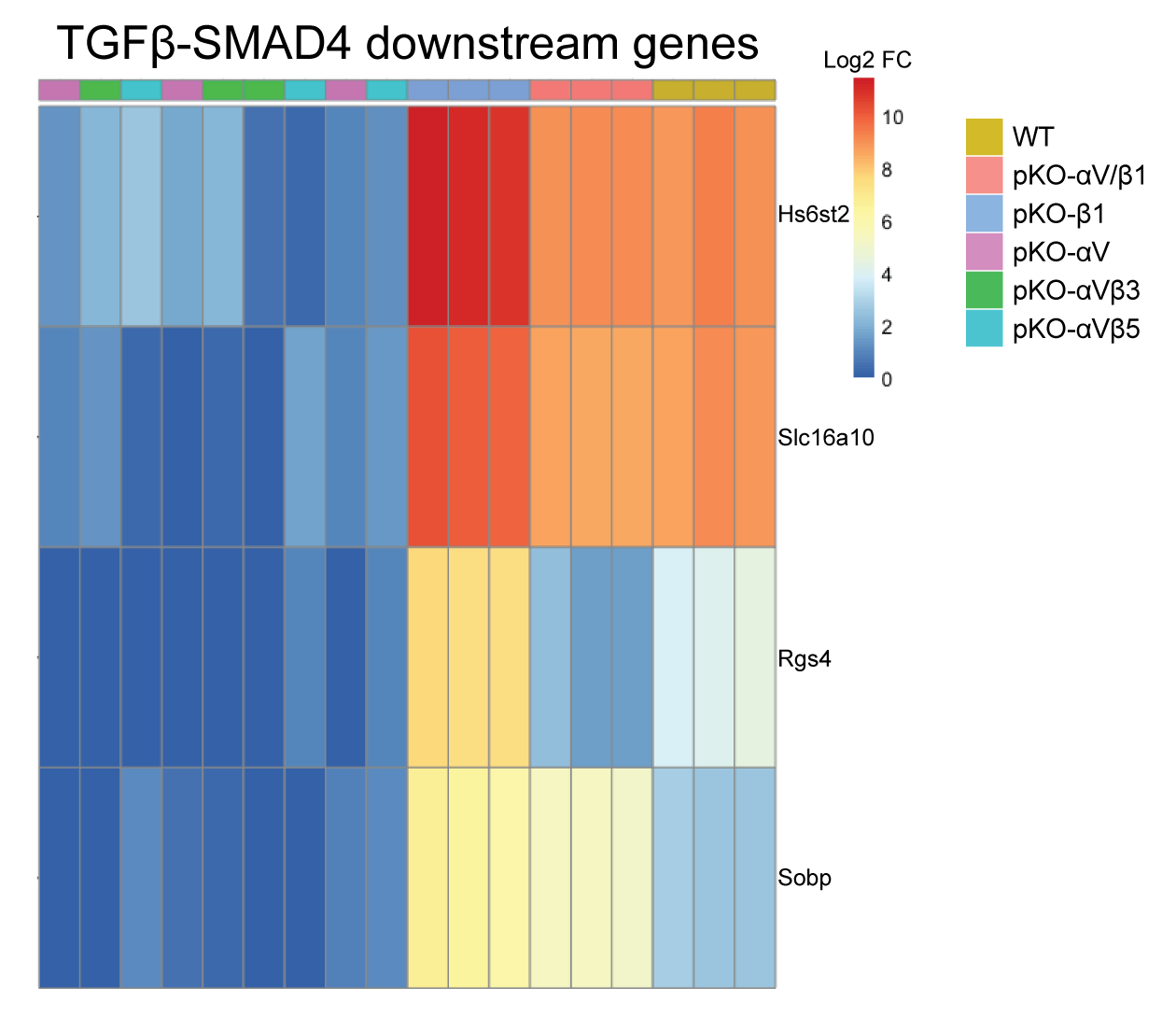

**Supplementary Figure 3. TGFβ-SMAD4-regulated genes show integrin-identity-dependent expression.** The heatmap of the different fibroblast lines depicts upregulation or downregulation of genes downstream of the transcription factor SMAD4 in an integrin class or integrin heterodimer specific manner. Color code of heatmap indicates log2 fold change (Log2 FC).

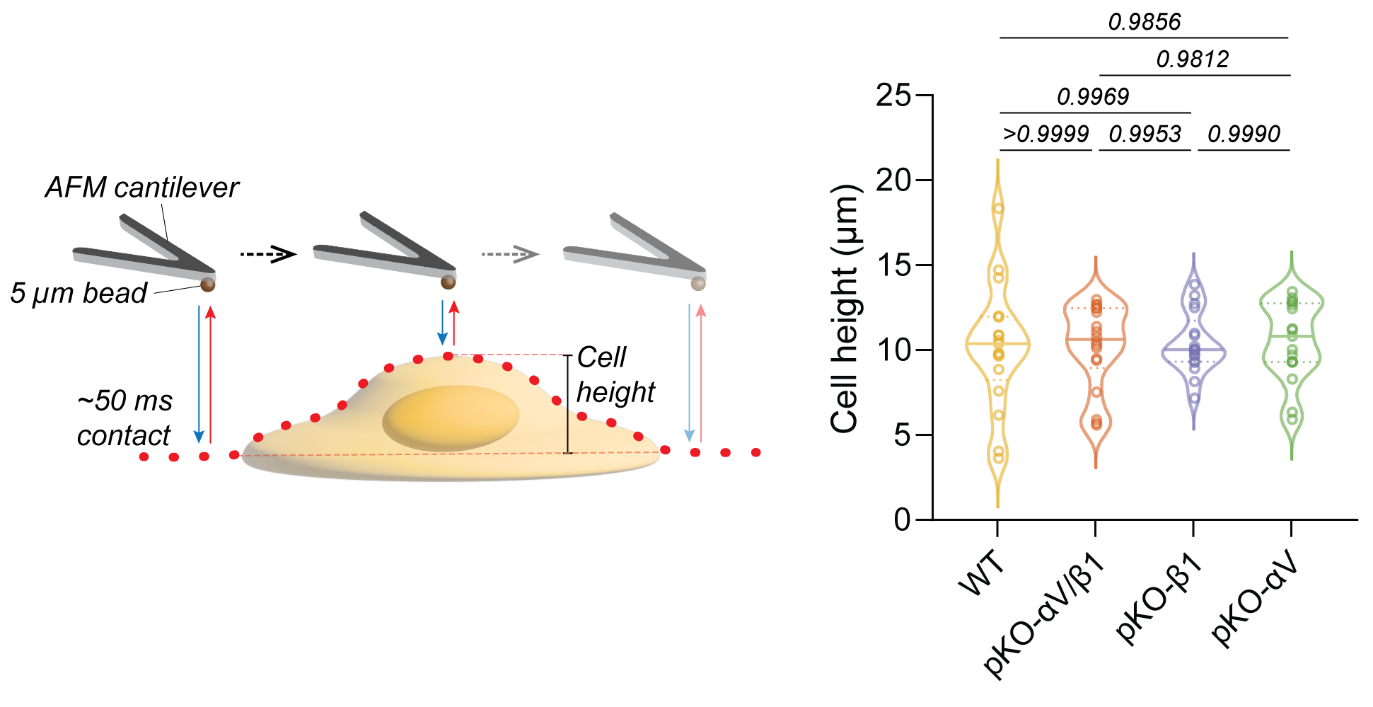
**Supplementary Figure 4. Cell heights of unperturbed fibroblast lines.** Overview of atomic force microscopy (AFM)-based cell height measurements using a 5 μm diameter bead glued to the free end of a microcantilever. The bead at the microcantilever was first used to approach the glass surface (reference) and then the cell surface above the nucleus at 5 μm s^–1^ until reaching a force of 500 pN. This procedure was repeated several times along a straight line spanning the glass surface and the fibroblast. Cell heights of WT (*n* = 17), pKO-αV/β1 (*n* = 18), pKO-β1 (*n* = 17), and pKO-αV (*n* = 17) fibroblasts after 1 h of being seeded on fibronectin-coated glass surfaces, and before being subjected to 20 or 5 μm confinement as described in the main manuscript. In the violin plots, each data point corresponds to a single fibroblast. The solid line represents the median, the dotted line indicates the interquartile range. The width of the violin plot represents the density of data points at that value with wider sections indicating higher density. *p*-values obtained from a one-way ANOVA analysis with Tukey’s multiple comparisons test with a single pooled variance.

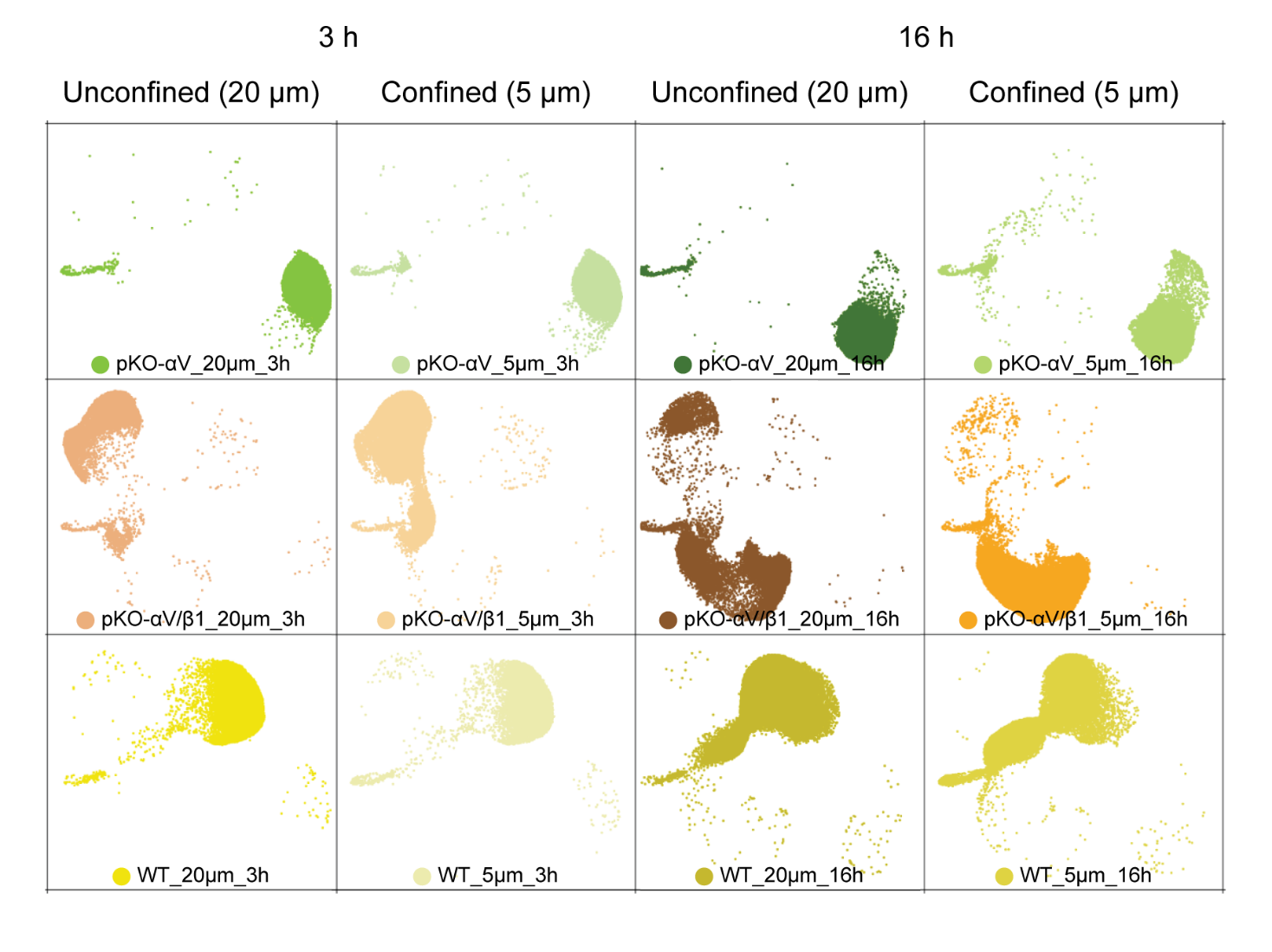
**Supplementary Figure 5. UMAP projections of transcriptomes of fibroblast lines unconfined and confined for different durations.** UMAP projections of pKO-αV, pKO-αV/β1, and WT fibroblasts being unconfined (20 µm) and confined (5 µm) for either 3 h or 16 h. Colors are based on the sample of origin. Each dot represents one single fibroblast. The data spread has been used to plot the dotted areas that define the transcriptomic regions of WT, pKO-αV/β1, and pKO-αV fibroblasts being unconfined and confined for 3 h or 16 h in Fig. 4b.

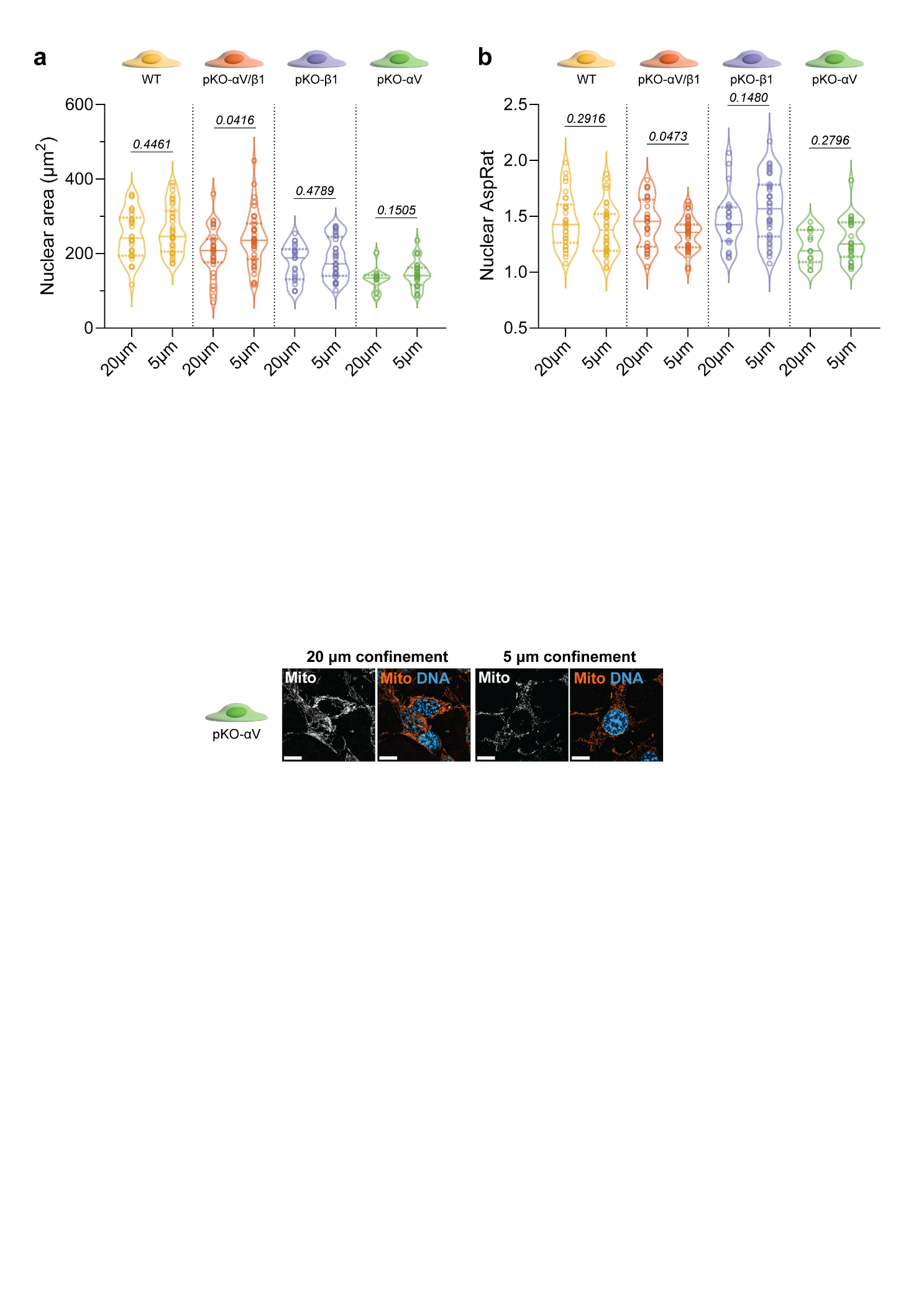

**Supplementary Figure 6. Nuclear characteristics of fibroblasts confined to different heights.** **a**, Nuclear area of unconfined (20 µm) and confined (5 µm) WT, pKO-αV/β1, pKO-β1, and pKO-αV fibroblasts. **b**, Nuclear aspect ratio (AspRat) of unconfined and confined WT [*n*(20 µm)= 30; *n*(5 µm)= 32], pKO-αV/β1 [*n*(20 µm)= 30; *n*(5 µm)= 32], pKO-β1 [*n*(20 µm)= 30; *n*(5 µm)= 32], and pKO-αV [*n*(20 µm)= 30; *n*(5 µm)= 32] fibroblasts. In the violin plots, each data point corresponds to a single fibroblast. The solid line represents the median, the dotted line indicates the interquartile range. The width of the violin plot represents the density of data points at that value with wider sections indicating higher density. *p*-values obtained from unpaired, two-sided, Mann-Whitney U-test.
